# Quantifying Local Perceptions of Environmental Change and Links to Community-Based Conservation Practices

**DOI:** 10.1101/2023.02.13.527316

**Authors:** Matt Clark, Haji Masoud Hamad, Jeffrey Andrews, Vicken Hillis, Monique Borgerhoff Mulder

**Affiliations:** Boise State University - Human Environment Systems; Revolutionary Government of Zanzibar Department of Forestry - Pemba; Max Planck Institute for Evolutionary Anthropology

**Keywords:** Community-based conservation, Mangroves, Environmental change, Participatory mapping

## Abstract

Approximately two billion people — a quarter of the earth’s population — directly harvest forest products to meet their daily needs. These individuals disproportionately experience the impacts of increasing climatic variability and global biodiversity loss, and must disproportionately alter their behaviors in response to these impacts. Much of the increasingly ambitious global conservation agenda relies on voluntary uptake of conservation behaviors in such populations. Thus, it is critical to understand how individuals in these communities perceive environmental change and use conservation practices as a tool to protect their well-being. To date however, there have been no quantitative studies of how individual perceptions of forest change and its causes shape real-world conservation behaviors in forest dependent populations. Here we use a novel participatory mapping activity to elicit spatially explicit perceptions of forest change and its drivers across 43 mangrove-dependent communities in Pemba, Tanzania. We show that perceptions of mangrove decline drive individuals to propose stricter limits on fuelwood harvests from community forests only if they believe that the resultant gains in mangrove cover will not be stolen by outsiders. Conversely, individuals who believe their community mangrove forests are at high risk of theft actually decrease their support for forest conservation in response to perceived forest decline. High rates of inter-group competition and mangrove loss are thus driving a ‘race to the bottom’ phenomenon in community forests in this system. This finding demonstrates a mechanism by which increasing environmental decline may cause communities to forgo conservation practices, rather than adopt them, as is often assumed in much community-based conservation planning. However, we also show that when effective boundaries are present, individuals are willing to limit their own harvests to stem such perceived decline.

## 1 Introduction

### 1.1 Problem statament

Diverse and healthy ecosystems are unequivocally our best insurance against the worsening impacts of climate change (Isbell et al. 2015; Loreau et al. 2001; Oliver et al. 2015; Lloret et al. 2012). Yet, increasingly intensive resource extraction from ecosystems over the last 150 years has greatly attenuated their ability to buffer human communities against impacts such as fires and flooding (Parks et al. 2016; Alongi 2008). Simultaneously, this switch from low to high intensity resource use has diminished global biodiversity on a magnitude only seen five other times in our planet’s history, further accelerating climate change (Caro et al. 2022).

Recent land use intensification strongly reflects the displacement of local communities and traditional practices by large-scale producers and outside economies (Stephens et al. 2019; Ellis et al. 2021; Bird et al. 2019). It is then largely recognized that effective and equitable conservation efforts must to empower local communities to set resource management priorities and design strategies to achieve them (Fernández-Llamazares et al. 2020; Garnett et al. 2018). Thus, achieving global conservation goals hinges, at least in part, on local community engagement and the decisions that individuals in those communities make in the environment (Gatiso et al. 2018).

Numerous studies have shown that resource users can reliably detect long-term changes in their local environments (Lauer and Aswani 2010; Early-Capistrán et al. 2022; Tengö and Belfrage 2004). However, it is still unclear how individuals perceptions’ of environmental change affect their choices to limit resource use, restore ecosystems, or otherwise change their behaviors (Paloniemi et al. 2018). In particular, as pointed out by Meyfroidt (2013), few studies have linked individual perceptions of threats and change in natural resources with observed conservation behaviors and preferences (although see Nyangoko et al. 2022). Further, a recent systematic review of 128 studies of voluntary adoption of conservation behaviors showed a dearth of research on the subject in non-Western populations (Thomas-Walters et al. 2022).

In her foundational work, Elinor Ostrom described a set of conditions that, when met, promote cooperative behaviors in natural resource management settings (Ostrom 1990). Among these conditions, Ostrom identifies the need to clearly demarcate and enforce proprietary access to group resources through physical and/or social boundaries (Ostrom 1990). Three decades of scrutiny via case studies and meta analyses from across the globe further cement this conclusion (Cox, Arnold, and Tomás 2010; Cox 2014; Cumming et al. 2020). In a recent set of theoretical models, Andrews and others (2022, 2023) delineate the social-ecological evolutionary mechanisms by which excluding outsiders promotes sustainable resource management behavior and cooperation in the face of threats to the local environment. However, the reverse is also true. These models show that in the absence of strong social or physical boundaries, perceived degradation of local resources may cause a ‘race to the bottom’ phenomenon where individuals are incentivized to extract all they can before the resource is gone (Andrews et al. 2023).

This theory explicitly predicts that environmental degradation should promote preferences for limiting resource extraction when theft from outsiders is low. And degradation should conversely promote preferences for increasing resource extraction when theft from outsiders is high, because the gains made by sustainable management may be eroded by outsiders and never realized by the local community (Andrews et al. 2023). This process has however not yet been examined empirically. An empirical test of these mechanisms is critical for building further theory in conservation science and for applying scientific insights to real-world resource management. For example, individuals make resource management decisions under the backdrop of past exposure to external conservation interventions and within a range of acceptable community norms (Hayes et al. 2022; Gómez-Baggethun and Ruiz-Pérez 2011). Thus, we must observe how theorized processes of behavioral change in response to environmental degradation operate in the real-world in order to have confidence in their general importance and applicability.

In this study, we perform an empirical test of how perceived environmental degradation and threat of resource theft from outsiders affect individuals’ conservation behaviors and preferences. We achieve this by implementing a novel participatory mapping activity to collect quantitative, spatially explicit perceptions of mangrove cover change in Pemba, Tanzania. We then link these perceptions of mangrove change with a questionnaire of individual perceptions of mangrove theft and self-reports of conservation behaviors and preferences. We specifically look at individuals’ self-reported frequency of patrolling behavior to protect community mangrove forests from outsiders and preferences for limits on the amount of fuelwood that community members can harvest from those forests. We assess these dynamics while simultaneously considering the impact that a major conservation initiative on the island (see section below) may have had on individuals’ conservation behaviors and preferences in the communities involved. We interpret the results of this analysis in light of their relation to theoretical work on the subject of perceived environmental change and resource boundary efficacy on conservation behaviors, thus increasing their generalizability and decreasing the probability of spurious findings (Smaldino and McElreath 2016).

### 1.2 Field site

This study examines community-based mangrove conservation in Pemba Island, Tanzania, the smaller of the two Zanzibari islands, identified as part of the Coastal Forests of Eastern Africa biodiversity hotspot. Like much of the developing world, Pemba has been subject to a series of conservation initiatives that stretch back to the colonial period, with novel initiatives increasing in frequency since the late 1990s. These begin with British colonial afforestation programs and the gazetting of forest reserves by both the British and post-revolutionary governments in the 1960s (Chachage 2000). Following 50 more years of initiatives driven by a number of Scandinavian countries, in 2010 the Reduced Emissions from Deforestation and Land Degradation program (REDD+) identified 18 wards (*shehia*) in Pemba as appropriate for piloting their payments for ecosystem services conservation framework (Burgess et al. 2010; RGZ 1996; Nations 1992). The REDD+ project intended to pay communities to forego harvesting fuelwood and timber and cease farm expansion inside of designated areas in each of the 18 selected shehia. The objective of this intervention was to slow deforestation, reduce greenhouse gas emissions, and reduce poverty. While hope for this project waxed and waned over several years among Pemban communities, these payments were never delivered and the 18 selected shehia ultimately showed no measurable benefit in forest cover (Andrews et al. 2021; Collins et al. 2022).

Alongside the proliferation and succession of these conservation projects, the population on the island has grown by approximately 2.9% each year (estimate from 2012 - 2022; more than triple the global average), increasing the need for the production of timber, fuelwood, and other forest products (URT 2023). Prior research shows that approximately 90% of rural Pemban households rely exclusively on forest products (fuelwood and charcoal) to meet their daily cooking needs (RGZ 2014; Ely et al. 2000). Further, these forest products account for 27% of total household income (Andrews and Borgerhoff Mulder 2022). This local need for forest products is driving a median deforestation rate of 3.4% per year in the forests of the island (Collins et al. 2022).

Many individuals across Pemba recognize that forests provide valuable ecosystem services such as erosion control, among many others. Thus, there is a conflict between the desire to safeguard local community forests, while still meeting daily needs. We find extensive evidence that individuals adapt to this challenge by stealing forest products from the community forests of other shehia; 70% of residents blamed their neighboring shehia for deforestation of their own local forests, and 31% of residents report having whole trees cut and stolen by outsiders (Borgerhoff Mulder, Caro, and Ngwali 2021 and unpublished data 2017).

Widespread cutting of mangroves in particular has caused considerable decline of mangrove cover and resultant flooding and saltwater intrusion in many mangrove adjacent communities (Andrews and Borgerhoff Mulder 2022). In response, many communities and community members therein have taken it upon themselves to prohibit outsiders from harvesting (or “stealing”) from their community mangroves and to reduce the harvests of their own community members. This generally takes the form of the establishment of village and shehia conservation committees, mangrove patrols to exclude outsiders from harvesting from community forests, mangrove planting, and setting specific fuelwood harvest limits. There is nevertheless considerable variability in preferences and practices of these actions on the island, both between and within shehia (Borgerhoff Mulder, Caro, and Ngwali 2021). We use this variability in individual preferences for limiting harvests and patrolling behavior as the outcomes of interest in this analysis.

## 2 Methods

### 2.1 Data collection

#### 2.1.1 Participatory mapping activity

We collected data on individual perceptions of environmental change using a participatory mapping methodology in order to elicit fine-scale, spatially explicit perceptions of change. This methodology builds on that of Herrmann et al. (2014) to tangibly link participant responses with specific locations and provide a more accurate measure than would be possible with a simple questionnaire (Emmel 2008; Cadag and Gaillard 2012). Over an eight-month field season in 2022, we were able to implement this methodology in 43 of the 49 shehia on the island which contain mangrove forest (Fig. 1). The six shehia not included in the study were excluded due to time and funding constraints, rather than for any systematic purpose. In each of these 43 shehia, we randomly selected five men and five women to participate in this activity, which resulted in a final sample size of 423 after dropping seven responses due to incomplete survey information.

**Figure 1:**
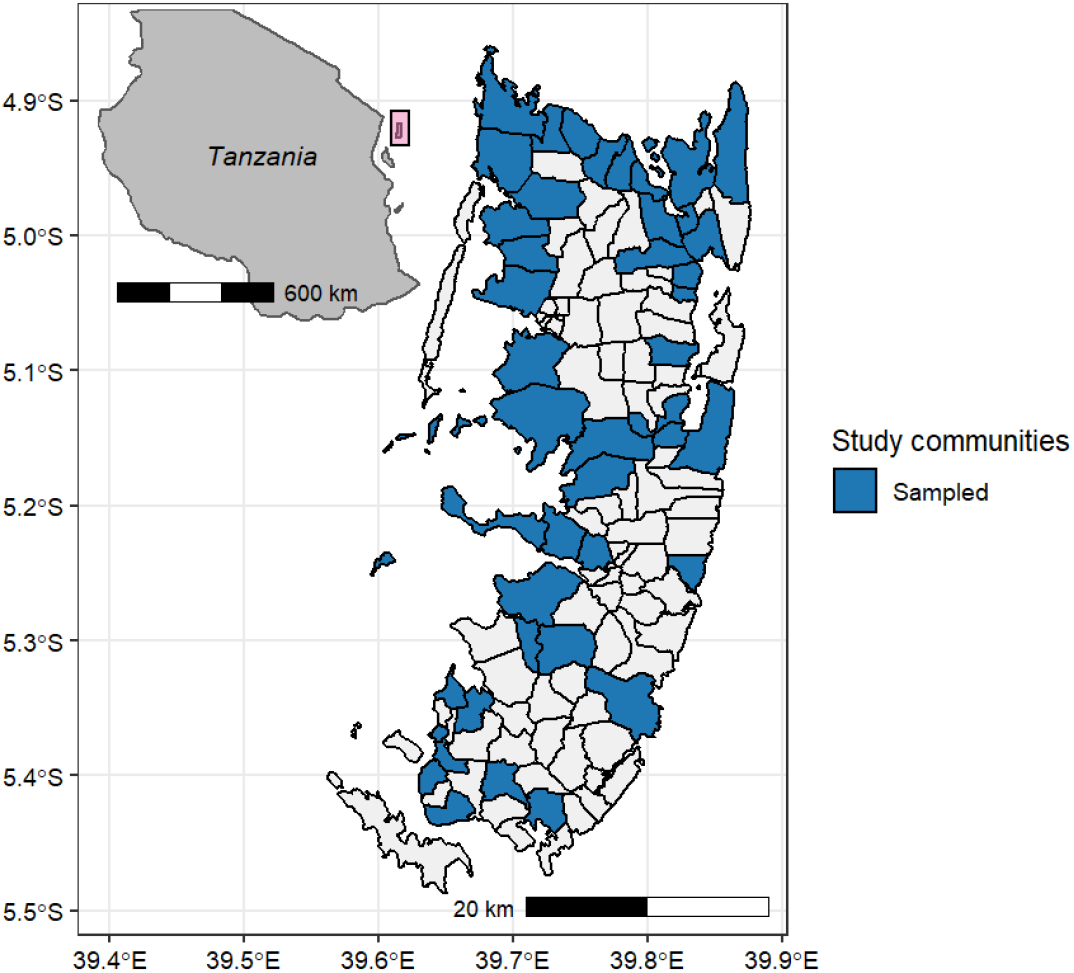
Wards (*shehia*) surveyed in this study. Large map shows the island of Pemba, Tanzania with each of the shehia where data were collected for this study shaded in blue. Inset map shows the location of Pemba in relation to the Tanzanian mainland.

The participatory mapping activity began with a workshop format where we established a shared understanding of our goals and did a simple mapping orientation, as most of the local population does not regularly use maps to navigate their environment. Each participant was then provided with a gridded basemap of their community, with towns, roads, bodies of water, cultural landmarks (e.g. mosques), and any protected areas labeled to help with orientation. Each grid cell corresponded to 0.5 km2 area. After a further orientation we asked participants to identify their own place of residence and other important locations to verify their basic understanding of the map. The final group task was to mark (initially with buttons until consensus was reached, then with a pen) each grid cell where mangrove forest is present. Thus, the workshop-style component of the participatory mapping activity ended once each participant was adequately oriented to a gridded map of their community, and each grid containing mangrove forest was marked identically across all participant maps (Fig. 2).

**Figure 2:**
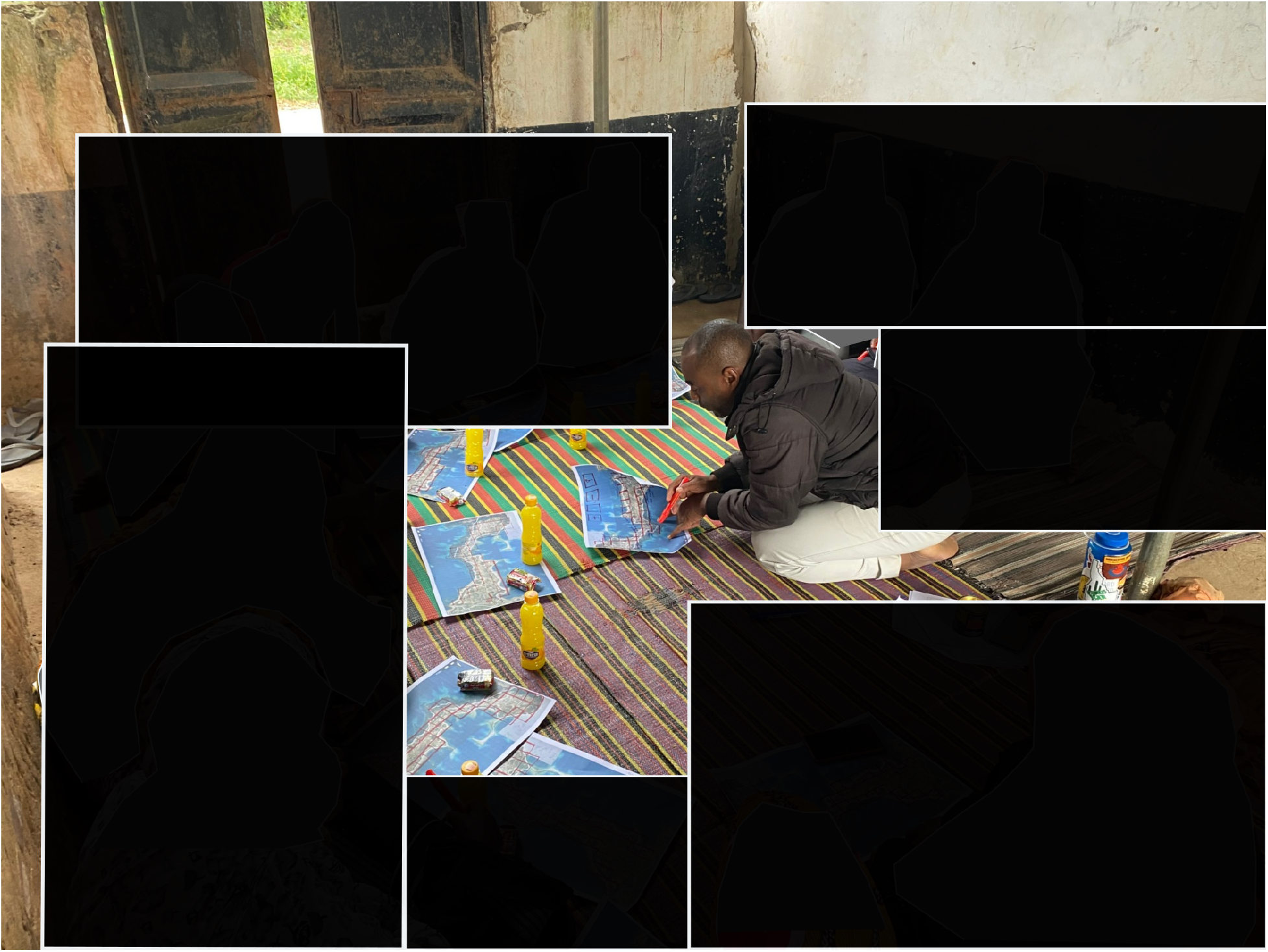
Example of the participatory mapping activity used in this study. Grid squares containing mangrove cover are first identified as a group, then respondents individually record their perceived change in each square. Photo shows H. M. Hamad explaining the individual response portion of the activity.

For the remainder of the participatory mapping activity and the questionnaire following, all participants responded individually. With the consensus map of mangrove locations inhand, each participant was asked to indicate, for each grid cell containing mangrove, whether they felt the tree cover in that area had increased, stayed the same, or decreased in the last year. Participants could also indicate that they were not sure about how mangrove cover had changed. An example of a completed map can be found in figure S1. The total number of grid cells in which a participant indicated that the mangrove cover had declined in the last year was tallied to produce an estimate of the perceived percent decline in community mangrove forest cover for each respondent.

#### 2.1.2 Questionnaire

Following the participatory mapping activity all participants completed an individual questionnaire with the help of research staff. The purpose of this questionnaire was to elicit responses regarding conservation behavior and preferences, perceived pressure of theft from outsiders, and general demographic information. Specifically, participants used a binary response to indicate whether or not they ever engage in patrols to protect community mangrove forests from theft from outsiders. If yes, participants listed the number of mangrove patrols that they estimated they had performed in the past month. Participants also indicated their preferences’ for harvest limits on themselves and other community members who rely on community mangroves to collect fuelwood. This outcome variable was collected as an integer value corresponding to the number of fuelwood bundles that they would like to limit themselves and their fellow community members to harvesting each month.

To quantify individuals’ perceptions of theft from outsiders in their community mangroves, we asked respondents to estimate the number of outsiders they believe come to their shehia to harvest fuelwood each week. We asked participants to provide their best guess of where these individuals generally come from in order to ensure they were describing individuals from outside their shehia, rather than a smaller village-level group. Finally, we recorded the gender and occupation of each participant through multiple choice questions and asked whether they were a member of a village or shehia conservation committee using a binary choice question. The full questionnaire instrument can be found in the supplemental material (S2).

### 2.2 Analysis

We performed two separate analyses in this research. The first (model 1; eq 1) was designed to estimate the effects of perceived decline of community mangroves and perceived mangrove theft on preferences for in-group harvest limits on fuelwood. In accordance with current best practices for causal inference, we constructed a directed acyclic graph to determine what parameters needed to be controlled for in order to estimate the direct effects of interest (Table 1 & Fig. S2a) (McElreath 2020; Westreich and Greenland 2013; Pearl 2009). In this process, we explicitly describe the complete hypothesized causal pathway between our predictors and outcome of interest and identify other variables, and associations between them, that may be affecting the outcome through separate causal paths (Pearl 2009). We then control for these alternative causal paths in order to capture accurate effect sizes for our direct effects of interest.

**Table 1:** List of variables and associated models.

| Variable | Collected from | Estimand or control | Model used in |
| --- | --- | --- | --- |
| **PREDICTORS** |  |  |  |
| Occupation | Questionnaire | Control | Model 1 (equation 1) |
| Perception of community mangrove change in the past year | Participatory mapping activity | Estimand | Both |
| Perceived number of outsiders stealing from community mangroves per week | Questionnaire | Estimand | Both |
| Interaction between perceived mangrove change and perceived mangrove theft | Questionnaire & participatory mapping activity | Estimand | Model 1 (equation 1) |
| Size of community mangrove area | Participatory mapping activity | Control | Both |
| Community included in REDD+ initiative | Previous research | Estimand | Both |
| Member of shehia or village conservation committee | Questionnaire | Estimand | Both |
| Gender | Questionnaire | Estimand | Model 2 (equation 3) |
| **OUTCOMES** |  |  |  |
| Preferred community fuelwood harvest limit | Questionnaire | Outcome | Model 1 (equation 1) |
| Number of mangrove patrols conducted in the past month | Questionnaire | Outcome | Model 2 (equations 2 & 3) |

We used a Poisson distributed generalized linear mixed-model operationalized in a Bayesian framework to estimate the direct effects of interest (estimands) (Table 1). We estimated the effect of the interaction between perceived mangrove loss and the theft pressure from outsiders on community mangrove forests. As identified using the directed acyclic graph, we controlled for participant occupation, the size of the community mangrove area, whether or not the participant was a member of a village or shehia conservation committee, and whether the shehia was one of the 18 exposed to the failed REDD+ intervention on the island. Finally, as we used a mixed model, we estimated a varying intercept (***β0**_j_*) for each of the 43 study shehia. This model is formalized in equation 1.

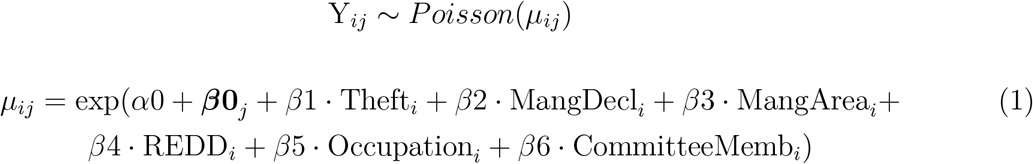

The secondary analysis for this research (model 2; eq 2 & 3) estimated the effects of perceived mangrove theft from outsiders and forest cover loss on reported respondent engagement in community mangrove patrols. To adequately model the data generating process for participation in community mangrove patrols, we operationalized this research question as a *hurdle* process (Zuur et al. 2009). In this framework, we model the joint outcome of whether or not a respondent is likely to report engaging in mangrove patrols at all (Bernouli distributed with probability *θ*) and if so, the number of patrols that they report engaging in each month (zero-truncated negative binomial distribution with mean *μ* and dispersion *ϕ*). Thus, the probability mass function is shown in equation 2.

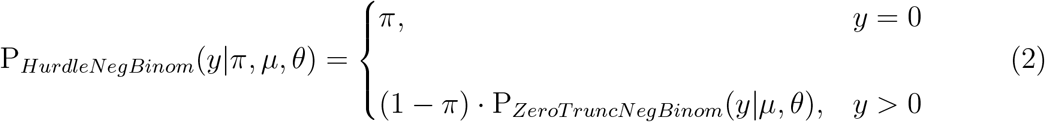

Again, for this analysis, we selected the parameter set necessary to estimate the direct effects of interest using a directed acyclic graph (S2b). Through this procedure, we concluded that to estimate the effect of perceived theft and forest loss on patrolling behavior, we must account for the size of the community mangrove area, the gender of the participant, whether or not the participant was a member of a village or shehia conservation committee, and whether the shehia was one of the 18 subjected to the failed REDD+ intervention on the island. In this model we substitute gender for participant occupation because gender affects both occupation and patrolling behavior, thus including both gender and occupation would result in estimating the effect of gender along two separate causal paths. In model 1 we do not assume that participant gender should affect their preferences for in-group harvest limits. We again used a Bayesian mixed-model, where we estimate a varying intercept for each of the 43 shehia in our study 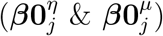. This model is formalized in equation 3.

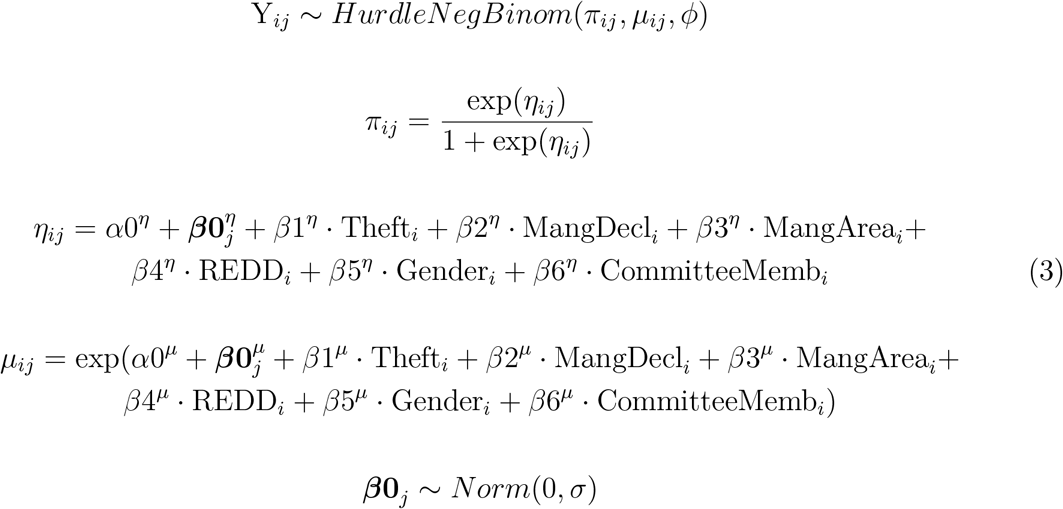

For both models, we used regularizing priors as recommended by Gelman et al. (2008) for producing conservative coefficient estimates. Both models exhibited adequate convergence of Markov chains, adequate posterior predictive capacity, and *R* values equal to 1 for all coefficients (S3). All data for this project and the R and STAN code used in these models is available in the Open Science section.

## 3 Results

As this is a Bayesian analysis, we consider any parameter estimate in which the inner 0.9 quantile of the posterior mass does not overlap zero to be statistically significant. This threshold is standard in the literature as it indicates that 95% or greater of the entire probability mass of sample estimates sit on one side of zero and therefore a 0.95 probability of a true effect given the data (Goodrich et al. 2020).

### 3.1 Preferences for fuelwood harvest limits

We find strong evidence that the interaction between individual perceptions of mangrove degradation and perceptions of mangrove theft from outsiders significantly affects preferences for fuelwood harvest limits from community mangroves. Two thousand draws from the posterior distribution indicated a 0.98 probability that the interaction term has a positive effect on the outcome (Fig. 3). In respondents who reported no perceived theft in their community mangrove forests, an increase in perceived mangrove decline from 0% to 50% of the community mangrove area resulted in an expected decrease in preferred harvest limits from 2.73 to 2.36 bundles of fuelwood. Respondents who reported that 100% of their community mangroves were declining in cover in turn reduced their fuelwood harvest limits to 2.04 bundles.

**Figure 3:**
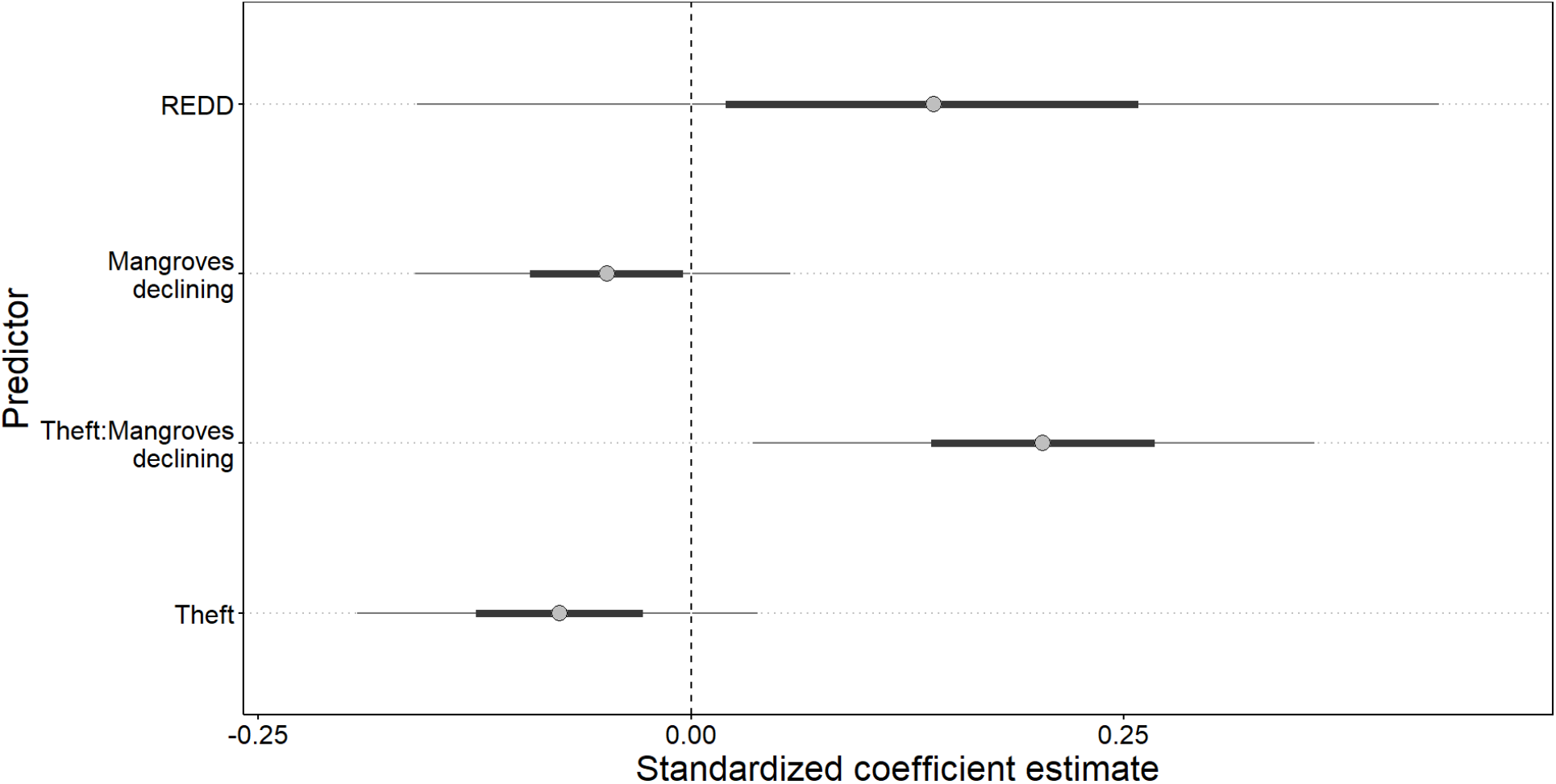
Standardized posterior estimates from the model shown in equation 1 used to estimate the drivers of preferences for limiting fuelwood use. The thick bars show the inner 50% of the posterior distribution and the thin bars show the inner 90% of the posterior distribution (credibility interval).

The interaction term indicates that this trend is reversed in individuals who perceive high levels of mangrove theft from outsiders. In these respondents, an increase in perceived mangrove decline from 0% to 50% of the community mangrove area resulted in a loosening of preferred harvest limits from 1.24 bundles of fuelwood to 2.73. Interestingly, the strength of this trend increased as more of the mangrove area was perceived as being in decline. Respondents who perceive the highest levels of theft and report that 100% of the community mangrove area is declining are expected to report a preference for a harvest limit of 6.07 bundles, a nearly five fold increase from those who perceive that 0% of the community mangrove area is in decline. The marginal effect of this interaction term, given a mean value of all other predictors, can be seen in figure 4.

**Figure 4:**
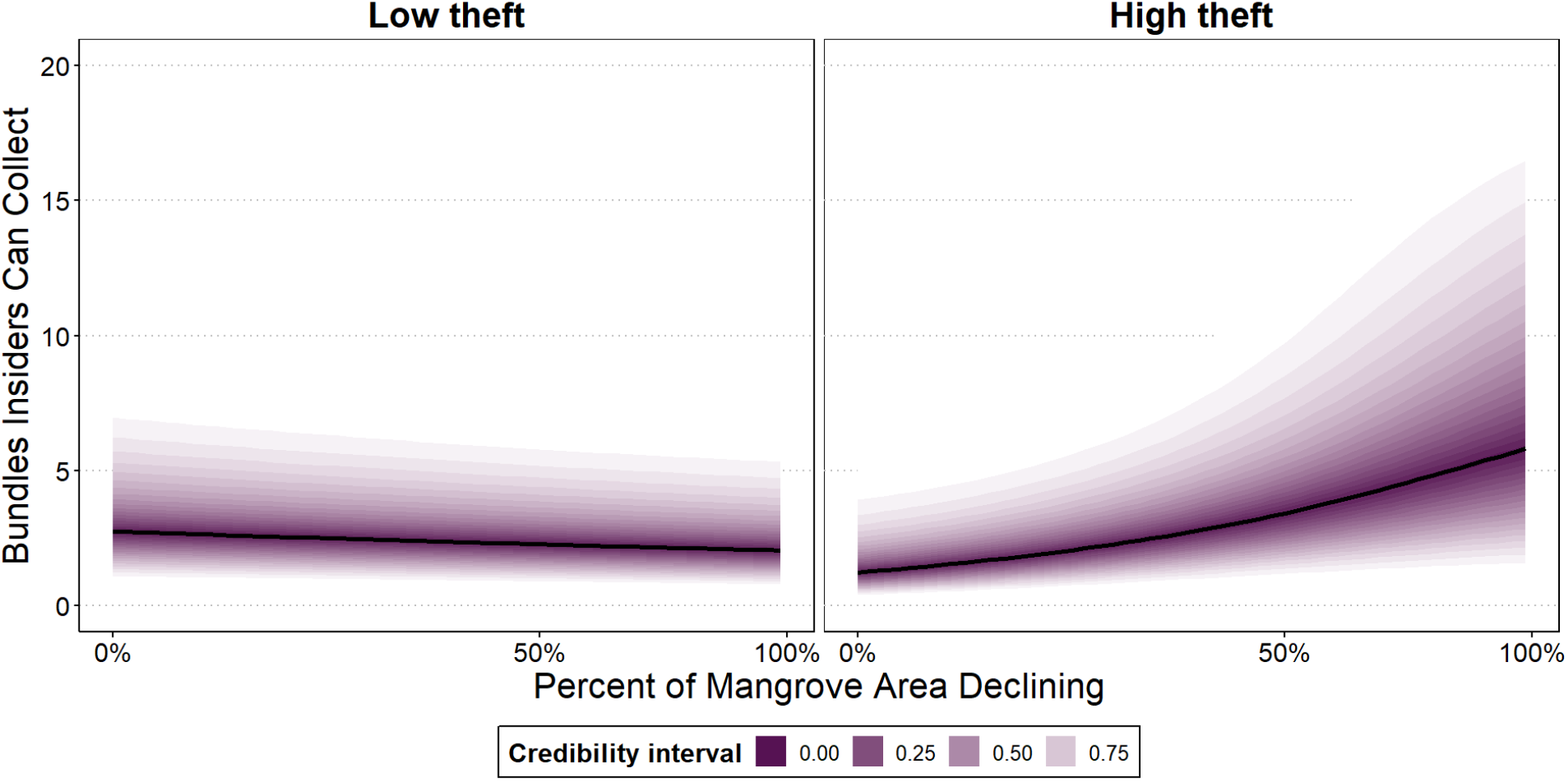
Marginal effect of the interaction of individual perception of mangrove decline and perceived intergroup theft on individual preference for in-group fuel-wood harvest limits. Low theft shows the effect of perceived mangrove decline when perceived theft was near 0 and high theft shows the effect when perceived theft was at the highest recorded value. The marginal effect shows the effect of these predictors at a mean value of all other predictors. Black lines show median model estimates. Shading those the credibility interval.

Finally, shehia who were part of the failed REDD+ initiative on the island showed a slight, although statistically insignificant, increase in preferred harvest limits compared to individuals in shehia where the REDD+ project was never introduced (Fig. 3). This effect is not statistically significant as the proportion of samples greater than zero is 0.79, representing a 0.79 probability of a true effect given our data.

### 3.2 Mangrove patrolling behavior

The coefficient estimates from the regression described in equation 3 show that patrolling behavior is likely driven by different processes than are preferences for restricting fuelwood harvests. The Bernoulli component of the model indicates that neither perceived mangrove theft or perceived mangrove decline significantly affected whether or not individuals reported engaging in mangrove patrols at all. The posterior distribution of the Bernoulli component of the model resulted in a statistically insignificant 0.87 probability that perceived theft increases the likelihood that individuals engage in mangrove patrols. Perceptions of mangrove decline had essentially no effect on likelihood of patrolling behavior (Fig. 5). Similarly, perceptions of mangrove decline and theft also had essentially no effect on the number of patrols an individual engaged in (Fig. 6).

**Figure 5:**
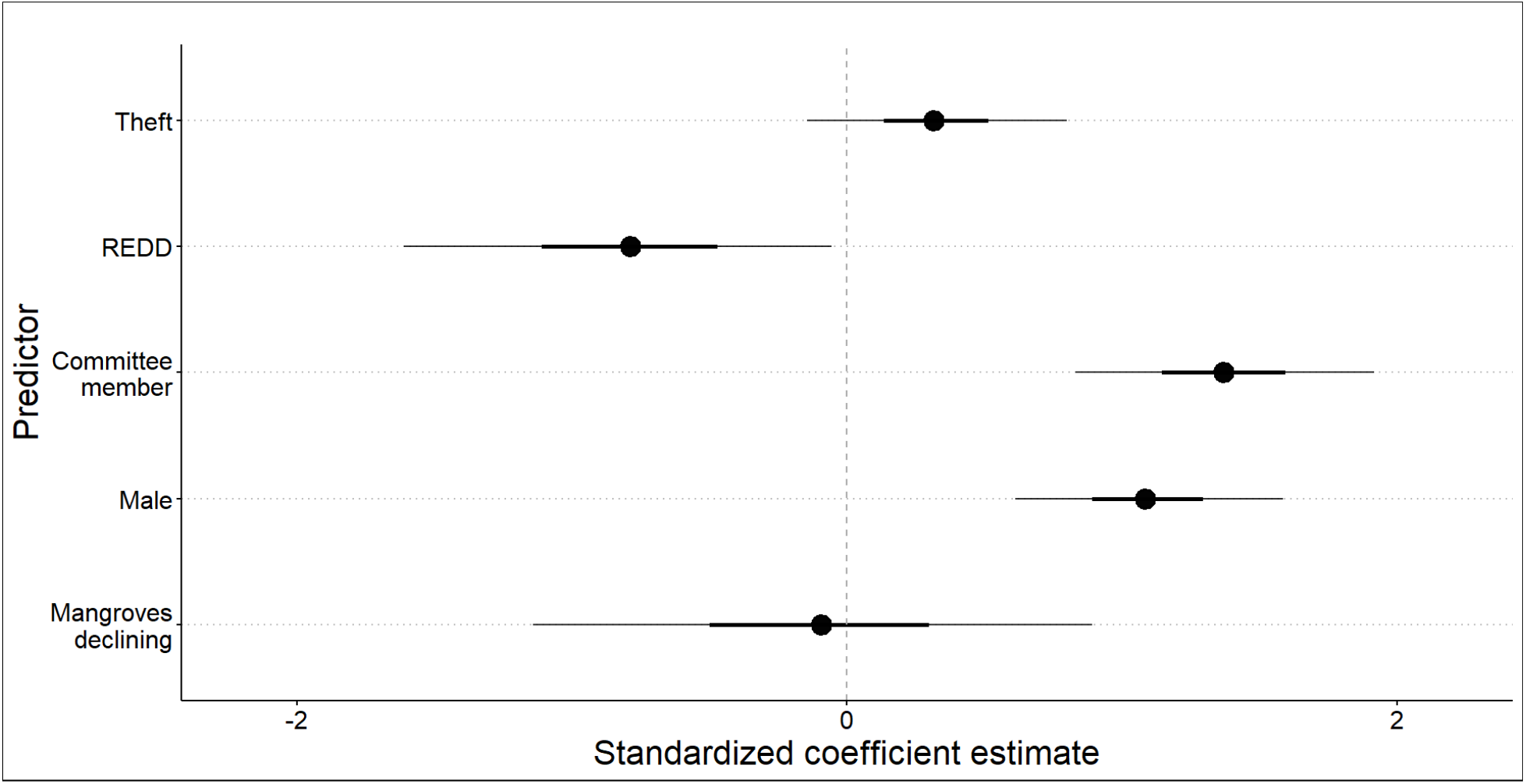
Standardized posterior estimates for the Bernoulli component of the model estimating the effect of these predictors on patrolling behavior (eq. 3). The Bernoulli component estimates the effect that the predictors have on whether or not individuals engage in patrolling behavior at all. The thick bars show the inner 50% of the posterior distribution and the thin bars show the inner 90% of the posterior distribution (credibility interval).

**Figure 6:**
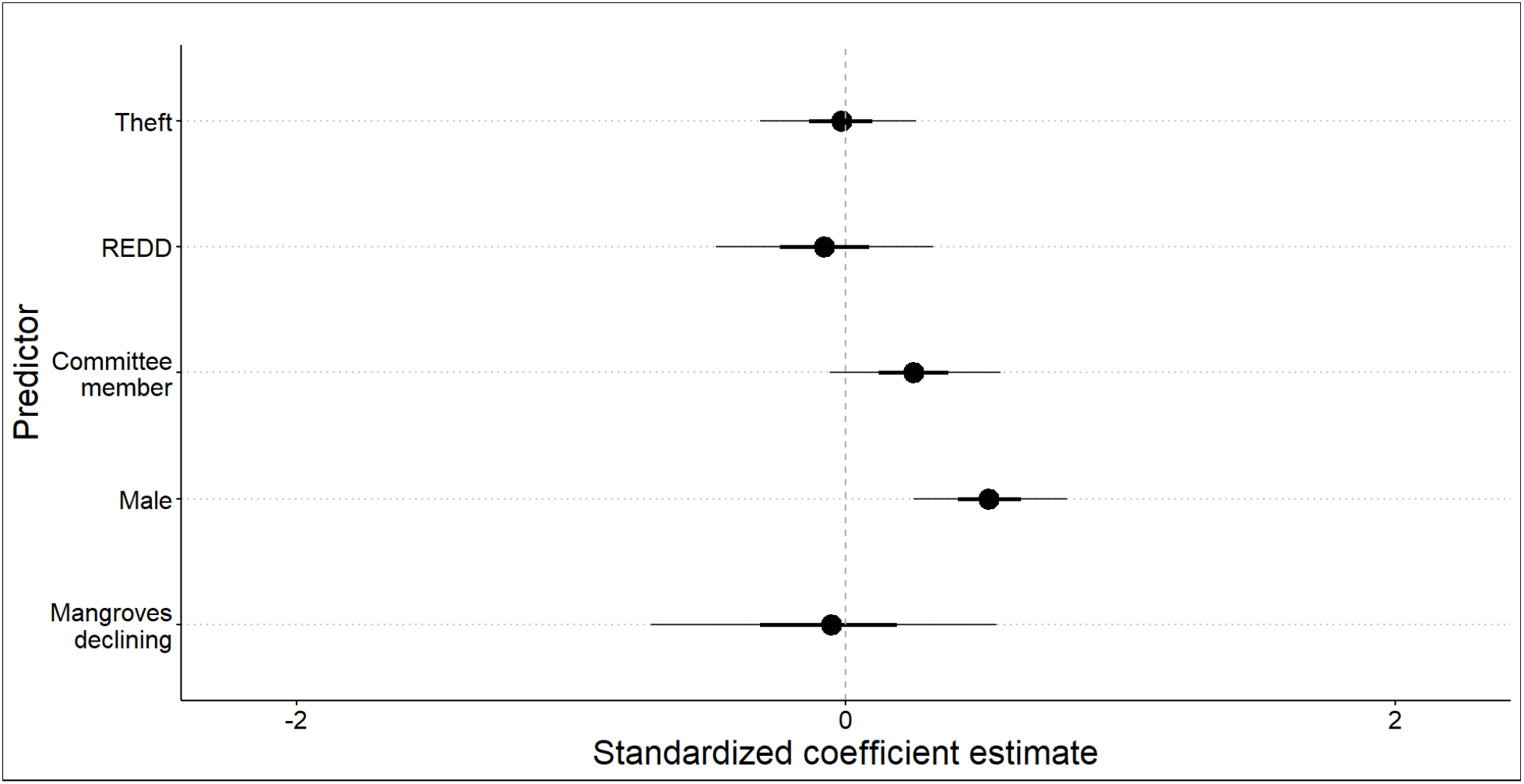
Standardized posterior estimates for the zero-truncated negative binomial component of the model estimating the effect of these predictors on patrolling behavior. The zero-truncated negative binomial component estimates the effect that the predictors have on the number of patrols that individuals engage in. The thick bars show the inner 50% of the posterior distribution and the thin bars show the inner 90% of the posterior distribution (credibility interval).

Men were significantly more likely than women to engage in patrols and to engage in a greater number of patrols (Fig. 5 & 6). Given a mean value for all other predictors, the probability that women reported engaging in patrols at all was 0.17 and the probability that men reported engaging in patrols at all was 0.38. Of men and women who reported patrolling, the median number of patrols performed by each gender in the last month was 6 and 3 respectively. Lastly, membership in a shehia or village conservation committee was significantly associated with individuals reporting going on patrols at all (Fig. 5), but was not significantly associated with the number of patrols they reported engaging in, as only 91% of model estimates were greater than 0 (Fig. 6).

The outputs of this model also indicate that past community exposure to REDD+ significantly decreased the probability of individual engagement in mangrove patrols. Specifically, there is a 0.96 probability that individuals from shehia selected for the failed REDD+ project were less likely to engage in mangrove patrols at all compared to those from shehia not exposed to the REDD+ project (Fig. 5). Given a mean value for all other predictors, the probability that individuals in shehia that were part of the REDD+ project reported engaging in patrols at all was 0.17, compared to a 0.32 probability for individuals from shehia not exposed to REDD+. However, this predictor was not significantly associated with the number of patrols that individuals engaged in (Fig. 6).

## 4 Discussion

### 4.1 Relation to & deviation from theory

In this research, we sought to test the theory that individual perception of environmental degradation will result in increased participation and support for conservation only if a lack of effective boundaries does not diminish the benefits of such conservation behaviors. Our data strongly support this intuition and furthermore show that perceived environmental degradation can actually decrease support for conservation if the threat of out-group freeriders is high. Thus, this finding, in combination with the theoretical development by Andrews et al. (2022, 2023), helps to detail the mechanisms underlying Ostrom’s first tenet that reliable boundaries are critical for sustainable common-pool resource management.

This research begins to fill the gap identified by Meyfroidt (2013), namely that little is known about how individuals use conservation behaviors to respond to perceived environmental change. These analyses reveal that different types of conservation behaviors are likely affected differently by perceived environmental change. While preferences for limiting resource use were greatly affected by perceptions of environmental change and its causes, behaviors to enforce resource boundaries were not. One reason for this result could be that individuals are hesitant to deviate from community norms. For example, women were much less likely to report engaging in mangrove patrols than men, even if they had identical perceptions of mangrove theft and decline, and a similar history with conservation programming. Additionally, patrolling is largely conducted by members of a village or shehia conservation committee; non-committee members were unlikely to begin engaging in this behavior solely of their own accord.

We speculate that because patrolling behavior is a visible action, pressure to adhere to local norms may operate more strongly on this outcome than on preferences for allowable community harvests which may be privately held. There is a growing body of literature on the adoption of conservation behaviors and scaling up of conservation projects to which this insight might be applicable (e.g. Mahajan et al. 2020; Mills et al. 2019; Clark, Andrews, and Hillis 2022). For example, theoretical models and analyses of empirical data may assume different social and ecological drivers of different classes of conservation actions. This field may then benefit by defining categories of conservation actions such as ‘in-group regulatory behaviors’ and ‘out-group exclusionary behaviors,’ or predominantly ‘environmentally-driven’ versus ‘socially-driven’ actions, among many other possible categorizations.

One interesting and somewhat unexpected important predictor emerged for both fuelwood harvest limits and mangrove patrolling. Past community exposure to the failed REDD+ project on the island was significantly associated with reduced probability of engagement in mangrove patrols, and showed a non-significant (p=0.79), yet interesting positive association with individual preferences for fuelwood harvest limits (these individuals preferred less stringent harvest limits). We hesitate to draw strong conclusions given these data, as this effect was not the primary question of the study (Tredennick et al. 2021). Yet, these trends are well aligned with theories regarding motivational crowding (Rode, Gómez-Baggethun, and Krause 2015; Frey and Jegen 2001). Along these lines, we speculate that past promises of payments for conservation behaviors, such as reducing fuelwood use and community forest patrols, may have crowded out individuals’ motivations to engage in such behaviors in the absence of payments (Cinner et al. 2021). Although there are other indications from a larger sample of individuals and broader environmental context (not limited to mangroves) collected in Pemba in 2017 that preferences for conservation did persist in communities exposed to the REDD+ intervention (Andrews and Borgerhoff Mulder 2023). This effect may then be nonlinear or context dependent.

### 4.2 Policy implications

As the negative impacts of climate change continue to affect communities of small-scale producers around the world, nature-based solutions, such as mangrove protection and restoration, are increasingly posited to buffer individuals against the worst impacts (E. Cohen-Shacham et al. 2016; Emmanuelle Cohen-Shacham et al. 2019). We show here that the uptake of nature-based solutions may be greatly hindered by a lack of clear social or physical boundaries to protect the benefits accrued by such actions. Yet, actions to exclude out-group members from community resources are costly. Our results show that they are so costly, that in fact, even when individuals perceive them as necessary, they will not perform them without some degree of social license (e.g. membership in a shehia conservation committee). Thus, this study suggests that support in the form of training and funding for community-based conservation initiatives specifically to demarcate and protect resource boundaries may increase their ability to combat the negative impacts of climate change through conservation. Such a policy may have dual benefits, directly stopping harvests from outsiders and supporting the endogenous emergence of sustainable in-group norms.

When gains from conservation behaviors are not eroded by outsiders, we show that individuals respond to perceived environmental degradation by supporting stricter limits on resource harvests. This result is promising for the prospect of meeting global conservation goals through community-based initiatives. The status of many resources are, however, not easily observable to local communities and even observed changes may be forgotten as individuals’ baselines for resource condition shift (Papworth et al. 2009). We emphasize then that supporting communities in effectively monitoring both local resources and the social benefits gained from protecting them is critical for the success of community-based conservation (Jones et al. 2013; Salerno et al. 2021).

### 4.3 Limitations and future work

The primary limitation in this research was non-random exclusion of the six shehia that we were unable to include due to time and funding constraints. However, our extensive ethnographic experience in Pemba does not lead us to believe that these shehia should fundamentally differ from those sampled in a way that would alter the results of this research. Specifically, these shehia do not greatly differ from those sampled in the importance of mangroves to the community, exposure to REDD+, or rates of environmental change. While we did not foresee the incomplete sampling of the 49 total shehia that contain mangrove forest at the onset of the data collection, the data collection scheme could have been improved by randomizing the order in which the shehia were visited for data collection.

Another key limitation of this work is that we rely on self-reported conservation preferences and behaviors for our outcomes of interest. The insights provided here would be bolstered if the realized conservation behaviors of participants could be observed. Future work might perform a similar participatory mapping activity with a random sample of a community after researchers host a tree planting activity or other conservation oriented event. Researchers may then record whether or not respondents attended the tree planting activity.

Conservation science would also benefit from a comprehensive examination of the effect that failed or terminated conservation projects, such as the REDD+ initiative on Pemba Island, have on local conservation preferences and behaviors (eg. Chervier, Le Velly, and Ezzine-de-Blas 2019; Massarella et al. 2018). Our results shown here are exploratory as this phenomenon was not the intended subject of study, but they may be an early signal of an important trend. Further, our measure of REDD+ exposure was at the community level, whether or not the shehia was one of the 18 selected for the intervention, and our outcomes were at the individual level. This finding would be strengthened by measuring individual exposure to REDD+ at the individual level as well.

## 5 Conclusion

In this paper we uncovered an important interaction between perceptions of environmental degradation and exposure to resource theft on two different types of conservation behaviors (harvest limits and community patrols). Put simply, individuals who are not exposed to theft while simultaneously experiencing resource decline are motivated to protect that dwindling resource. In contrast, individuals who *are* exposed to high levels of theft while simultaneously experiencing resource decline are motivated to actually *weaken* harvest limits, presumably in a race to grab what they can while it’s still available.

We also show that perceived mangrove degradation and theft from outsiders do not significantly affect individual engagement in patrols to exclude outsiders from stealing mangroves from community forests. Instead, this behavior is performed only by specific members of the community. Thus, as theft increases between communities, there is little mechanism to reduce it. And as theft is left largely unregulated, the ‘race to the bottom’ phenomenon causes in-group members to also harvest rapidly from community forests.

This social-ecological mechanism highlights the importance of clearly defined boundaries detailed by Ostrom in her first principle (1990). This research then echoes the importance of clear and effective boundaries and enforcement in community-based conservation efforts, and the positive endogenous changes in self-regulation that can follow in the wake of stronger boundaries.

## 6 Open Science

All code and data used in this project can be found at the Github link here: https://github.com/matthewclark1223/ParticipatoryMappingProj

## 7 Acknowledgements

We thank all of the community members who gave their time to help with this project and all of the community leaders (*sheha*) who helped coordinate our data collection efforts. We thank all personnel at the Pemba Department of Forests for ongoing support of our research. We also thank the Max Planck Institute for Evolutionary Anthropology Department of Human Behavior, Ecology and Culture for ongoing support of data collection and analysis.

## Notes

https://github.com/matthewclark1223/ParticipatoryMappingProj

### Competing Interest Statement

The authors have declared no competing interest.

## References

10 Alongi, Daniel M. 2008. “Mangrove Forests: Resilience, Protection from Tsunamis, and Responses to Global Climate Change.” Estuarine, Coastal and Shelf Science 76 (1): 1–13.

Andrews, Jeffrey, and Monique Borgerhoff Mulder. 2022. “Forest Income and Livelihoods on Pemba: A Quantitative Ethnography.” World Development 153 (May): 105817. https://doi.org/10.1016/j.worlddev.2022.105817.

Andrews, Jeffrey, and Monique Borgerhoff Mulder. 2023. “The Value of Failure: The Effect of an Expired Conservation Program on Residents’ Willingness for Future Participation.” Manuscript in review. Ecological Economics.

Andrews, Jeffrey, Monique Borgerhoff Mulder, Vicken Hillis, and Matthew Clark. 2022. “Adaptive Responses to Inter-Group Competition Over Natural Resources: The Case of Leakage, with Evidence from Pemba Tanzania.” {SSRN} {Scholarly} {Paper}. Rochester, NY. https://doi.org/10.2139/ssrn.4154871.

Andrews, Jeffrey, Tim Caro, Said Juma Ali, Amy C. Collins, Bidawa Bakari Hamadi, Hassan Sellieman Khamis, Abdi Mzee, Assaa Sharif Ngwali, and Monique Borgerhoff Mulder. 2021. “Does REDD+ Have a Chance? Implications from Pemba, Tanzania.” Oryx, 1–7. https://doi.org/10.1017/S0030605319001376.

Andrews, Jeffrey, Matthew Clark, Vicken Hillis, and Monique Borgerhoff Mulder. 2023. “Evolving Ostrom’s First Principle: The Dynamic Evolution of Common Property Rights and Their Role in Resource Management.” Research Square Preprint. https://doi.org/10.21203/rs.3.rs-2225599/v1.

Bird, Douglas W., Rebecca Bliege Bird, Brian F. Codding, and David W. Zeanah. 2019. “Variability in the Organization and Size of Hunter-Gatherer Groups: Foragers Do Not Live in Small-Scale Societies.” Journal of Human Evolution 131 (June): 96–108. https://doi.org/10.1016/j.jhevol.2019.03.005.

Borgerhoff Mulder, Monique, Tim Caro, and Assa Sharif Ngwali. 2021. “A Silver Lining to REDD: Institutional Growth Despite Programmatic Failure.” Conservation Science and Practice 3 (1): e312. https://doi.org/10.1111/csp2.312.

Burgess, Neil D., Bruno Bahane, Tim Clairs, Finn Danielsen, Søren Dalsgaard, Mikkel Funder, Niklas Hagelberg, et al. 2010. “Getting Ready for REDD+ in Tanzania: A Case Study of Progress and Challenges.” Oryx 44 (3): 339–51. https://doi.org/10.1017/S0030605310000554.

Cadag, Jake Rom D., and J. C. Gaillard. 2012. “Integrating Knowledge and Actions in Disaster Risk Reduction: The Contribution of Participatory Mapping.” Area 44 (1): 100–109. https://doi.org/10.1111/j.1475-4762.2011.01065.x.

Caro, Tim, Zeke Rowe, Joel Berger, Philippa Wholey, and Andrew Dobson. 2022. “An Inconvenient Misconception: Climate Change Is Not the Principal Driver of Biodiversity Loss.” Conservation Letters 15 (3): e12868. https://doi.org/10.1111/conl.12868.

Chachage, Chachage Seithy L. 2000. Environment, Aid and Politics in Zanzibar. Dar es Salaam University Press.

Chervier, Colas, Gwenolé Le Velly, and Driss Ezzine-de-Blas. 2019. “When the Implementation of Payments for Biodiversity Conservation Leads to Motivation Crowding-Out: A Case Study From the Cardamoms Forests, Cambodia.” Ecological Economics 156 (February): 499–510. https://doi.org/10.1016/j.ecolecon.2017.03.018.

Cinner, Joshua E., Michele L. Barnes, Georgina G. Gurney, Stewart Lockie, and Cristian Rojas. 2021. “Markets and the Crowding Out of Conservation-Relevant Behavior.” Conservation Biology 35 (3): 816–23. https://doi.org/10.1111/cobi.13606.

Clark, Matt, Jeffrey Andrews, and Vicken Hillis. 2022. “A Quantitative Application of Diffusion of Innovations for Modeling the Spread of Conservation Behaviors.” Ecological Modelling 473 (November): 110145. https://doi.org/10.1016/j.ecolmodel.2022.110145.

Cohen-Shacham, Emmanuelle, Angela Andrade, James Dalton, Nigel Dudley, Mike Jones, Chetan Kumar, Stewart Maginnis, et al. 2019. “Core Principles for Successfully Implementing and Upscaling Nature-Based Solutions.” Environmental Science & Policy 98 (August): 20–29. https://doi.org/10.1016/j.envsci.2019.04.014.

Cohen-Shacham, E., G. Walters, C. Janzen, and S. Maginnis, eds. 2016. Nature-Based Solutions to Address Global Societal Challenges. IUCN International Union for Conservation of Nature. https://doi.org/10.2305/IUCN.CH.2016.13.en.

Collins, Amy C., Mark N. Grote, Tim Caro, Aniruddha Ghosh, James Thorne, Jonathan Salerno, and Monique Borgerhoff Mulder. 2022. “How Community Forest Management Performs When REDD$\mathplus$ Payments Fail.” Environmental Research Letters 17 (3): 034019. https://doi.org/10.1088/1748-9326/ac4b54.

Cox, Michael. 2014. “Understanding Large Social-Ecological Systems: Introducing the SESMAD Project.” International Journal of the Commons 8 (2).

Cox, Michael, Gwen Arnold, and Sergio Villamayor Tomás. 2010. “A Review of Design Principles for Community-Based Natural Resource Management.” Ecology and Society 15 (4). https://www.jstor.org/stable/26268233.

Cumming, G. S., G. Epstein, J. M. Anderies, C. I. Apetrei, J. Baggio, Ö. Bodin, S. Chawla, et al. 2020. “Advancing Understanding of Natural Resource Governance: A Post-Ostrom Research Agenda.” Current Opinion in Environmental Sustainability, Resilience and complexity:Frameworks and models to capture social-ecological interactions, 44 (June): 26–34. https://doi.org/10.1016/j.cosust.2020.02.005.

Early-Capistrán, Michelle María, Elena Solana-Arellano, F. Alberto Abreu-Grobois, Gerardo Garibay-Melo, Jeffrey A. Seminoff, Andrea Sáenz-Arroyo, and Nemer E. Narchi. 2022. “Integrating Local Ecological Knowledge, Ecological Monitoring, and Computer Simulation to Evaluate Conservation Outcomes.” Conservation Letters n/a (n/a): e12921. https://doi.org/10.1111/conl.12921.

Ellis, Erle C., Nicolas Gauthier, Kees Klein Goldewijk, Rebecca Bliege Bird, Nicole Boivin, Sandra Díaz, Dorian Q. Fuller, et al. 2021. “People Have Shaped Most of Terrestrial Nature for at Least 12,000 Years.” Proceedings of the National Academy of Sciences 118 (17). https://doi.org/10.1073/pnas.2023483118.

Ely, Adrian V, AB Omar, AU Basha, SA Fakih, R Wild, et al. 2000. “A Participatory Study of the Wood Harvesting Industry of Charawe and Ukongoroni, United Republic of Tanzania.” A Participatory Study of the Wood Harvesting Industry of Charawe and Ukongoroni, United Republic of Tanzania.

Emmel, Nick. 2008. “Participatory Mapping: An Innovative Sociological Method.” Working {Paper}. Real Life Methods. http://eprints.ncrm.ac.uk/540/.

Fernández-Llamazares, Álvaro, Julien Terraube, Michael C. Gavin, Aili Pyhälä, Sacha M. O. Siani, Mar Cabeza, and Eduardo S. Brondizio. 2020. “Reframing the Wilderness Concept Can Bolster Collaborative Conservation.” Trends in Ecology & Evolution 35 (9): 750–53. https://doi.org/10.1016/j.tree.2020.06.005.

Frey, Bruno S., and Reto Jegen. 2001. “Motivation Crowding Theory.” Journal of Economic Surveys 15 (5): 589–611. https://doi.org/10.1111/1467-6419.00150.

Garnett, Stephen T., Neil D. Burgess, Julia E. Fa, Álvaro Fernández-Llamazares, Zsolt Molnár, Cathy J. Robinson, James E. M. Watson, et al. 2018. “A Spatial Overview of the Global Importance of Indigenous Lands for Conservation.” Nature Sustainability 1 (7): 369–74. https://doi.org/10.1038/s41893-018-0100-6.

Gatiso, Tsegaye T., Björn Vollan, Ruppert Vimal, and Hjalmar S. Kühl. 2018. “If Possible, Incentivize Individuals Not Groups: Evidence from Lab-in-the-Field Experiments on Forest Conservation in Rural Uganda: Individual Versus Community Incentives.” Conservation Letters 11 (1): e12387. https://doi.org/10.1111/conl.12387.

Gelman, Andrew, Aleks Jakulin, Maria Grazia Pittau, and Yu-Sung Su. 2008. “A Weakly Informative Default Prior Distribution for Logistic and Other Regression Models.” The Annals of Applied Statistics 2 (4): 1360–83. https://doi.org/10.1214/08-AOAS191.

Gómez-Baggethun, Erik, and Manuel Ruiz-Pérez. 2011. “Economic Valuation and the Commodification of Ecosystem Services.” Progress in Physical Geography: Earth and Environment 35 (5): 613–28. https://doi.org/10.1177/0309133311421708.

Goodrich, Ben, Jonah Gabry, Imad Ali, and Sam Brilleman. 2020. “Rstanarm: Bayesian Applied Regression Modeling via Stan.” R Package Version 2 (1).

Hayes, Tanya, Felipe Murtinho, Hendrik Wolff, María Fernanda López-Sandoval, and Joel Salazar. 2022. “Effectiveness of Payment for Ecosystem Services After Loss and Uncertainty of Compensation.” Nature Sustainability 5 (1): 81–88. https://doi.org/10.1038/s41893-021-00804-5.

Herrmann, Stefanie M., Ibrahima Sall, and Oumar Sy. 2014. “People and Pixels in the Sahel: A Study Linking Coarse-Resolution Remote Sensing Observations to Land Users’ Perceptions of Their Changing Environment in Senegal.” Ecology and Society 19 (3). https://www.jstor.org/stable/26269611.

Isbell, Forest, Dylan Craven, John Connolly, Michel Loreau, Bernhard Schmid, Carl Beierkuhnlein, T Martijn Bezemer, et al. 2015. “Biodiversity Increases the Resistance of Ecosystem Productivity to Climate Extremes.” Nature 526 (7574): 574–77.

Jones, Julia P. G., Gregory P. Asner, Stuart H. M. Butchart, and K. Ullas Karanth. 2013. “The ‘Why,’ ‘What’ and ‘How’ of Monitoring for Conservation.” In Key Topics in Conservation Biology 2, 327–43. John Wiley & Sons, Ltd. https://doi.org/10.1002/9781118520178.ch18.

Lauer, Matthew, and Shankar Aswani. 2010. “Indigenous Knowledge and Long-Term Ecological Change: Detection, Interpretation, and Responses to Changing Ecological Conditions in Pacific Island Communities.” Environmental Management 45 (5): 985–97. https://doi.org/10.1007/s00267-010-9471-9.

Lloret, Francisco, Adrian Escudero, Jose Maria Iriondo, Jordi Martinez-Vilalta, and Fernando Valladares. 2012. “Extreme Climatic Events and Vegetation: The Role of Stabi-lizing Processes.” Global Change Biology 18 (3): 797–805.

Loreau, Michel, Shahid Naeem, Pablo Inchausti, Jan Bengtsson, JP Grime, Andrew Hector, DU Hooper, et al. 2001. “Biodiversity and Ecosystem Functioning: Current Knowledge and Future Challenges.” Science 294 (5543): 804–8.

Mahajan, Shauna L., Arundhati Jagadish, Louise Glew, Gabby Ahmadia, Hannah Becker, Robert Y. Fidler, Lena Jeha, et al. 2020. “A Theory-based Framework for Understanding the Establishment, Persistence, and Diffusion of Community-based Conservation.” Conservation Science and Practice, October. https://doi.org/10.1111/csp2.299.

Massarella, Kate, Susannah M. Sallu, Jonathan E. Ensor, and Rob Marchant. 2018. “REDD+, Hype, Hope and Disappointment: The Dynamics of Expectations in Conservation and Development Pilot Projects.” World Development 109 (C): 375–85. https://ideas.repec.org//a/eee/wdevel/v109y2018icp375-385.html.

McElreath, Richard. 2020. Statistical Rethinking: A Bayesian Course with Examples in R and STAN. 2nd ed. New York: Chapman; Hall/CRC. https://doi.org/10.1201/9780429029608.

Meyfroidt, Patrick. 2013. “Environmental Cognitions, Land Change, and Social–Ecological Feedbacks: An Overview.” Journal of Land Use Science 8 (3): 341–67. https://doi.org/10.1080/1747423X.2012.667452.

Mills, Morena, Michael Bode, Michael B. Mascia, Rebecca Weeks, Stefan Gelcich, Nigel Dudley, Hugh Govan, et al. 2019. “How Conservation Initiatives Go to Scale.” Nature Sustainability 2 (10): 935–40. https://doi.org/10.1038/s41893-019-0384-1.

Nations, United. 1992. “Rio Declaration on Environment and Development.”

Nyangoko, Baraka P., Håkan Berg, Mwita M. Mangora, Mwanahija S. Shalli, and Martin Gullström. 2022. “Community Perceptions of Climate Change and Ecosystem-Based Adaptation in the Mangrove Ecosystem of the Rufiji Delta, Tanzania.” Climate and Development 14 (10): 896–908. https://doi.org/10.1080/17565529.2021.2022449.

Oliver, Tom H, Matthew S Heard, Nick JB Isaac, David B Roy, Deborah Procter, Felix Eigenbrod, Rob Freckleton, et al. 2015. “Biodiversity and Resilience of Ecosystem Functions.” Trends in Ecology & Evolution 30 (11): 673–84.

Ostrom, Elinor. 1990. Governing the Commons: The Evolution of Institutions for Collective Action. Cambridge University Press.

Paloniemi, Riikka, Teppo Hujala, Salla Rantala, Annika Harlio, Anna Salomaa, Eeva Primmer, Sari Pynnönen, and Anni Arponen. 2018. “Integrating Social and Ecological Knowledge for Targeting Voluntary Biodiversity Conservation.” Conservation Letters 11 (1): e12340. https://doi.org/10.1111/conl.12340.

Papworth, S.k., J. Rist, L. Coad, and E.j. Milner-Gulland. 2009. “Evidence for Shifting Baseline Syndrome in Conservation.” Conservation Letters 2 (2): 93–100. https://doi.org/10.1111/j.1755-263X.2009.00049.x.

Parks, Sean A., Carol Miller, John T. Abatzoglou, Lisa M. Holsinger, Marc-André Parisien, and Solomon Z. Dobrowski. 2016. “How Will Climate Change Affect Wildland Fire Severity in the Western US?” Environmental Research Letters 11 (3): 035002. https://doi.org/10.1088/1748-9326/11/3/035002.

Pearl, Judea. 2009. Causality. Cambridge University Press.

RGZ. 1996. “The Forest Management and Conservation Act No. 10 of 1996.” Legal Supplement to the Zanzibar Government Gazette.

RGZ. 2014. “Zanzibar’s Climate Change Strategy.”

Rode, Julian, Erik Gómez-Baggethun, and Torsten Krause. 2015. “Motivation Crowding by Economic Incentives in Conservation Policy: A Review of the Empirical Evidence.” Ecological Economics 117 (September): 270–82. https://doi.org/10.1016/j.ecolecon.2014.11.019.

Salerno, Jonathan, Chelsie Romulo, Kathleen A Galvin, Jeremy Brooks, Patricia Mupeta-Muyamwa, and Louise Glew. 2021. “Adaptation and Evolution of Institutions and Governance in Community-Based Conservation.” Conservation Science and Practice 3 (1): e355. https://doi.org/10.1111/csp2.355.

Smaldino, Paul E., and Richard McElreath. 2016. “The Natural Selection of Bad Science.” Royal Society Open Science 3 (9): 160384. https://doi.org/10.1098/rsos.160384.

Stephens, Lucas, Dorian Fuller, Nicole Boivin, Torben Rick, Nicolas Gauthier, Andrea Kay, Ben Marwick, et al. 2019. “Archaeological Assessment Reveals Earth’s Early Transformation Through Land Use.” Science 365 (6456): 897–902. https://doi.org/10.1126/science.aax1192.

Tengö, Maria, and Kristina Belfrage. 2004. “Local Management Practices for Dealing with Change and Uncertainty: A Cross-Scale Comparison of Cases in Sweden and Tanzania.” Ecology and Society 9 (3). https://www.jstor.org/stable/26267678.

Thomas-Walters, Laura, Jamie McCallum, Ryan Montgomery, Claire Petros, Anita K. Y. Wan, and Diogo Veríssimo. 2022. “Systematic Review of Conservation Interventions to Promote Voluntary Behavior Change.” Conservation Biology n/a (n/a): e14000. https://doi.org/10.1111/cobi.14000.

Tredennick, Andrew T., Giles Hooker, Stephen P. Ellner, and Peter B. Adler. 2021. “A Practical Guide to Selecting Models for Exploration, Inference, and Prediction in Ecology.” Ecology n/a (n/a): e03336. https://doi.org/10.1002/ecy.3336.

URT, Tanzania. 2023. “Tanzania Population and Housing Census 2022.” Population Distribution by Administrative Areas. https://www.nbs.go.tz/nbs/takwimu/Census2022/matokeomwanzooktoba2022.pdf.

Westreich, Daniel, and Sander Greenland. 2013. “The Table 2 Fallacy: Presenting and Interpreting Confounder and Modifier Coefficients.” American Journal of Epidemiology 177 (4): 292–98. https://doi.org/10.1093/aje/kws412.

Zuur, Alain F., Elena N. Ieno, Neil J. Walker, Anatoly A. Saveliev, and Graham M. Smith. 2009. “Zero-Truncated and Zero-Inflated Models for Count Data.” In Mixed Effects Models and Extensions in Ecology with R, edited by Alain F. Zuur, Elena N. Ieno, Neil Walker, Anatoly A. Saveliev, and Graham M. Smith, 261–93. Statistics for Biology and Health. New York, NY: Springer. https://doi.org/10.1007/978-0-387-87458-6_11.

