## Appendix S1 for "Quantifying Local Perceptions of Environmental Change and Links to Community-Based Conservation Practices"

**Appendix S1 – Participatory mapping example**

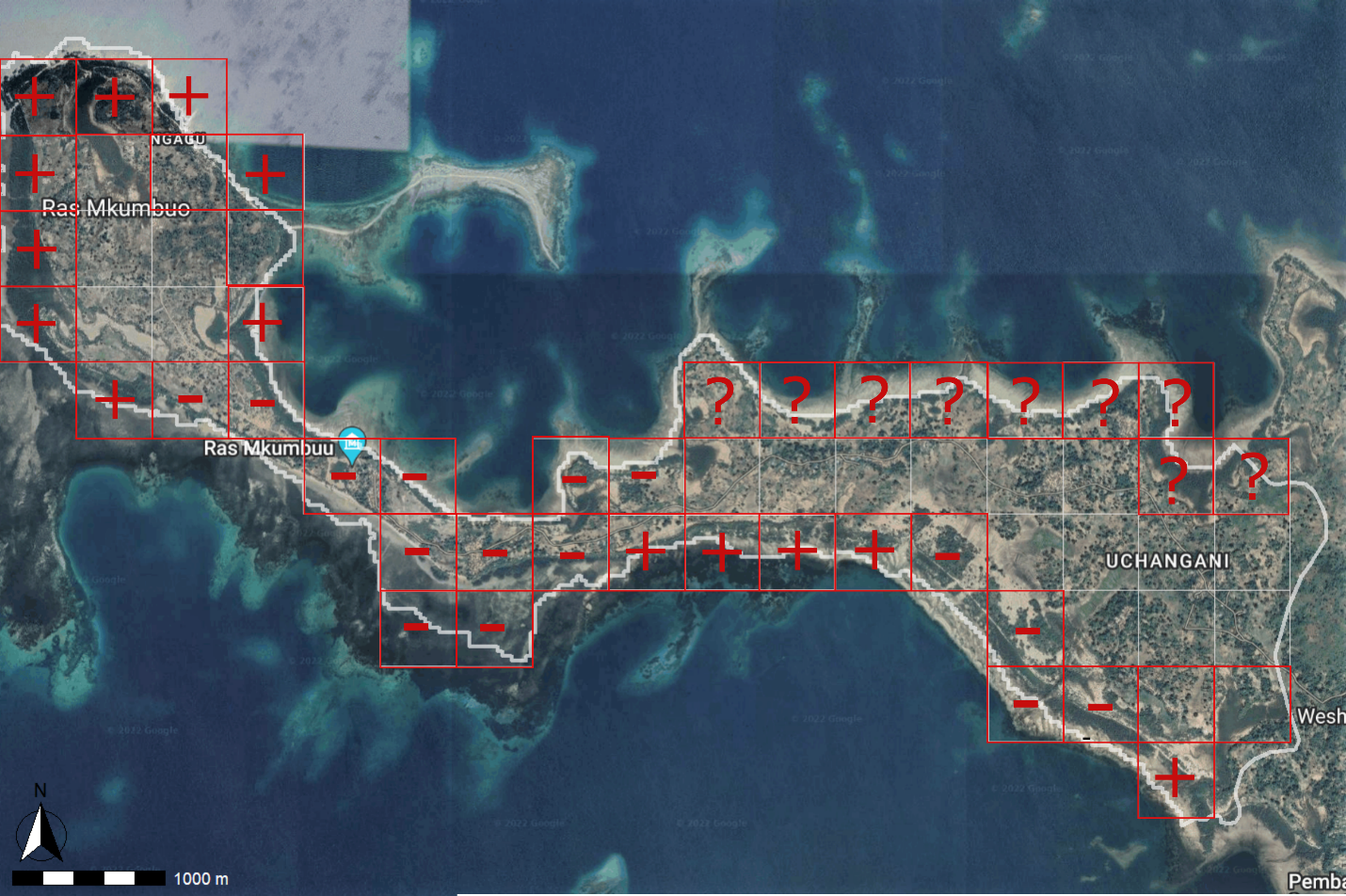

Figure S1: Stylized example of a completed map used in the participatory mapping activity in this research. Grid cells correspond to 0.5km^2^ areas. Outlined grid cells indicate community consensus on the presence of mangrove trees in the area. Plus (+) signs indicate individual perception that mangrove cover has increased in the area in the past year. Minus (-) signs indicate individual perception that mangrove cover has decreased in the area in the past year. Empty squares indicate individual perception of no change in mangrove cover in the past year. Question marks (?) indicate that the respondent does not know how mangrove cover in that specific area has changed in the past year.
