## Appendix S3 for "Quantifying Local Perceptions of Environmental Change and Links to Community-Based Conservation Practices"

**Appendix S3 – Model 2 posterior predictive check**

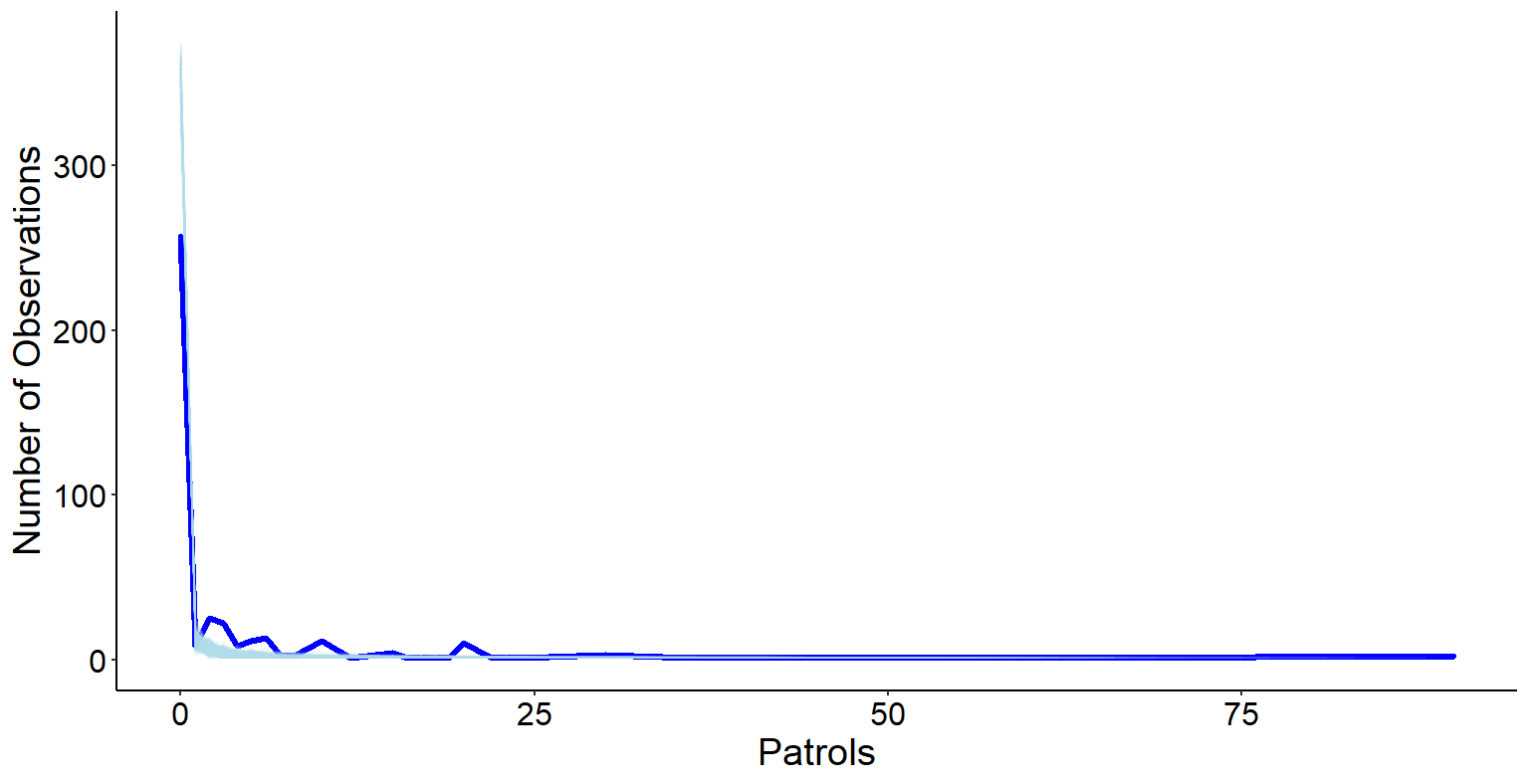

Figure S3: Posterior predictive check for model 2 used in this research. Dark blue line shows the true number of observed patrols. Light blue lines show the 2,000 draws from the model. We show that our model reliably reproduces data that match our observed data, further indicating adequate model fit in addition to adequate mixing of chains, rhat values equal to 1 for all model parameters, and lack of divergent transitions after warmup.
